# Brain Region-Specific Oligodendrocyte States Highlight Mitochondrial Gene Upregulation and Loss of Canonical Identity Signatures in Alzheimer’s Disease

**DOI:** 10.64898/2026.01.14.699226

**Authors:** Francesca Drummer, Nadia Dorosti, Charlene Hurler, Eduardo Beltrán, Louis B. Kümmerle, Fabian Theis, Sarah Jäkel

## Abstract

Oligodendrocyte dysfunction and heterogeneity are emerging as key contributors to Alzheimer’s disease (AD) pathology, though cellular mechanisms remain unclear. Using single-nucleus RNA sequencing across three cortical regions from the same individuals, we identified region- and disease-specific transcriptomic changes, including increased mitochondrial genes and loss of cell type signatures. These results highlight the importance of biologically-informed quality control and underscore the value of multi-regional analysis for understanding AD pathogenesis and progression.

## Main text

Emerging evidence has identified oligodendrocytes (OL) as a noteworthy cell type in the etiology of Alzheimer’s disease (AD)^1^. Although, thus far, transcriptomic studies have pointed to oligodendroglial dysfunction in AD, the pathomechanisms leading to their dysfunction and driving the progression of the disease across different cortical regions remain largely elusive.

To investigate brain region-specific transcriptomic changes of OL in AD, we performed single nucleus RNA-Sequencing (snRNA-seq) on post-mortem human brain tissue of six AD patients and age- and sex-matched controls each with an APOE ε3/ε3 background. Cortical grey matter samples partially depleted of neurons were collected from three areas per donor: prefrontal, temporal, and visual cortices (PFC, TC and VC, respectively). These regions were chosen to represent a snapshot of the temporal progression of the disease with the VC serving as a minimally affected internal control^2^ (Fig. 1A, Extended Data Table 1). Unlike previous studies, the present approach analyzes differently affected regions from the same donor, enabling cellular comparisons across disease progression, which has so far not been performed, especially in regard to OL.

**Figure 1:**
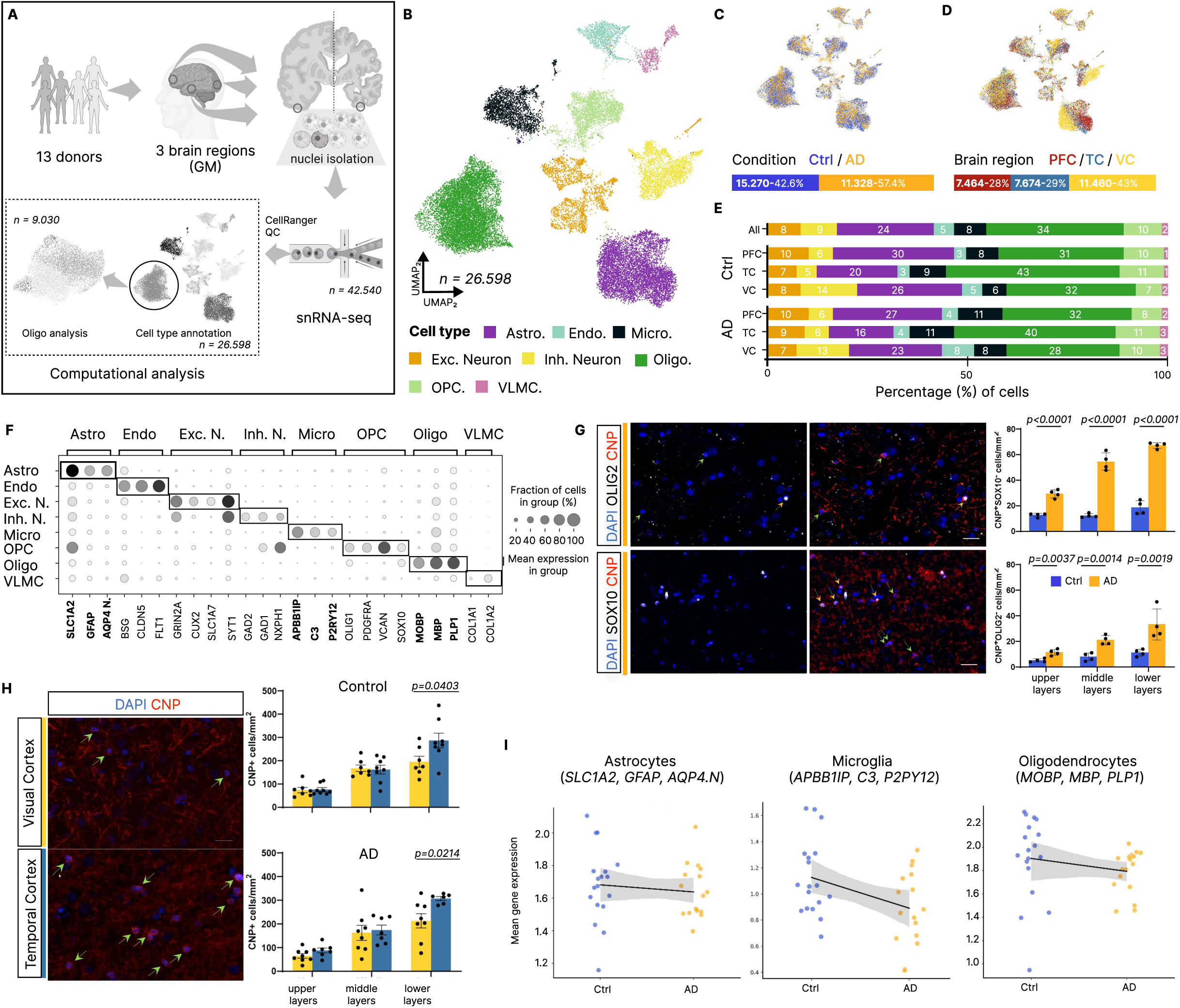
Cell type specific transcriptional profiles diverge across brain areas in AD pathology. **A** Study design showing snRNA-seq analysis of 13 donors across 3 brain regions (prefrontal, temporal and visual cortex) with nuclei isolation and sequential computational analysis of oligodendrocytes. **B-D** UMAP visualizations of all cell types after QC (n = 26,598 nuclei), with major cell populations (**B**), conditions - AD vs Control - (**C**) and cortical areas (**D**). **E** Percentage distribution of cellular composition across conditions and cortical areas (PFC, TC, VC). **F** Dotplot of selected marker genes for broad cell type annotation. **G** Representative immunofluorescence images of AD patient brains and quantifications of CNP+ OLIG2-/SOX10-oligodendrocytes in human brain tissue (n = 4 individuals per group), demonstrating a decrease of canonical marker expression in oligodendrocytes in AD. Scale bar = 50µm. **H** Quantifications of CNP+ oligodendrocytes (green arrows) across cortical layers in the visual and temporal cortices in control and AD brains (n = 7-8 individuals per group). Scale bar= 20µm. **I** Mean expression of canonical marker genes for astrocytes, microglia and oligodendrocytes between control and AD, demonstrating a general loss in AD.

After quality control (Extended Data Fig. 1, Extended Data Table 2), the remaining 26.598 nuclei clustered into seven major cell types by the expression of established canonical marker genes: OL (9.030 nuclei, *OLIG2*, *SOX10*), oligodendrocyte precursor cells (OPC, 2.530 nuclei, *PDGFRA*, *VCAN*), microglia (2.262 nuclei, *APBB1IP*, *C3*), astrocytes (6.420 nuclei, *SLC1A2*, *GFAP*), subtypes of both excitatory (2.377 nuclei, *CUX2*, *SLC1A7*) and inhibitory neurons (2.221 nuclei, *GAD1*, *GAD2*), endothelial cells (EC, 1240 nuclei, *CLDN5*, *FLT1*) and vascular leptomeningeal cells (VLMC, 518 nuclei, *COL1A1*, *COL1A2*) (Fig. 1B-F). Similarly to another study^3^ we observed mixed lineage clusters (Extended Data Fig. 2A-D, Extended Data Table 3). Although these might confer biological function, as demonstrated by co-expression of CD31 and CNP in some cells in the human brain (Extended Data Fig. 2E) we removed them in our analysis to focus on established OL clusters. Whilst disease condition did not influence proportional cluster abundance, assignment to a specific brain region did (Fig. 1E), likely due to different cellular architectures and functional specializations^4,5^.

In accordance with other studies^6–8^, we did not observe significant changes in cell type proportions in AD compared to control donors within the same region (Fig. 1E). When shifting our focus to OLs, contrary to what is commonly demonstrated by histological analysis^9,10^, we did not capture a loss of these cells in the disease condition, which was surprising as tissue-based microscopy approaches are better in determining cell frequencies. In fact, whilst most cell types remained stable across brain regions, we found a slight increase in OL proportions in the PFC and TC. To validate OL numbers and to identify whether there are further discrepancies between bioinformatic and histological analyses, we chose to immunohistochemically assess OL lineage cells (OLIG2) and mature OL (CNP) and EC (CD31) (Fig. 1G,H, Extended Data Fig. 3A,B) in a different patient cohort. Whilst OLIG2 and CNP+ OL were overall not changed between conditions, lower layers of the TC exhibited significantly higher numbers of mature OL, which supports our bioinformatic analysis and a recent live-imaging study that reported different OL densities in different cortical areas in rodents^11^. CD31+ EC however, showed a clear trend towards being increased in AD, which were not captured by our bioinformatic analysis. In brief, these data emphasize the importance of performing immunohistochemical validations to corroborate bioinformatic findings, and highlight compositional differences between different cortical regions.

Although OLIG2 numbers were not significantly changed in our analysis in AD, we observed weaker OLIG2 signals within cells during our analysis, raising the hypothesis that canonical marker expression and therefore the characteristic cell type signature might be lost in disease. Indeed, further validations using OLIG2/SOX10 and CNP confirmed a significant increase in the number of SOX10- or OLIG2- CNPase+ OL (Fig. 1G). We then quantified the expression of canonical marker genes for each cell type used for cluster annotation (Fig. 1F) and found a lower mean expression for OL, as well as for some other cell types across and within brain areas in AD (Fig. 1I, Extended Data Fig. 3C, Extended Data Table 4). Canonical marker genes were especially reduced in OLs in the TC and VC, which was quite unexpected, as the VC is generally expected not to be heavily impacted by the pathology^2^. Therefore, our bioinformatic data support the hypothesis of changed cell type signatures with decreased canonical marker gene expression. This finding might explain the discordance in the number of OL in AD between histopathological and transcriptional studies, as cluster annotation in snRNA-seq data does not solely rely on a single canonical cell marker.

Interestingly, we observed an increase in mitochondrial (MT) gene fraction over all cells with increasing Braak stages (Fig. 2A). Uniform MT gene fraction filtering is classically used as a quality control (QC) criterion^12^, however a recent study^13^ demonstrated a significant higher baseline MT gene fraction in cancer vs healthy cells. Therefore adaptive^14^ or nuclear fraction thresholding, such as MALAT1^15^ has been suggested as an alternative QC assessment. After confirming less variation of MALAT1 as compared to MT-fraction, across Braak stage in our data (Fig. 2A,B, Extended Data Fig. 4A), we applied biologically informed thresholding (Suppl. Tab 2), driven by condition and region (Extended Data Table 5), showing that MALAT1, MT-thresholding and other QC criteria consider different cells as low quality (Extended Fig. 4B). Subsequently, we still observed a high overall *MALAT1* expression across cells, but cell type specific proportions of low (<0.15) and high MT-fractions (>0.15) (Fig. 2C), with excitatory neurons and OL being on the higher end, and astrocytes and endothelial cells on the lower end. Further analysing MT-fraction across cortical areas in our data (Fig 2B) and Gabitto et. al, 2024^16^ (Extended Data Fig. 4D), we found an increase from PC to TC to VC, which we also observed in our Visium spatial transcriptomics data (Fig. 2D, Extended Data Fig. 5). If elevated MT-fraction represented poor sample quality, brain regions within the same donor would show a similar MT-fraction level. As the VC is affected later and less heavily by AD pathological burden than the PFC and TC, we would expect higher MT-fractions in PFC and TC due to a higher abundance of stressed and dying cells. Hence, our data demonstrate a condition-specific upregulation of MT-genes. Moreover, the MT-gene fraction along all Braak stages in another dataset (Extended Data Fig 4C) also suggests an upregulation in the earlier stages of the disease (Braak stage 3). Cellular stress has been reported to tether mitochondria to the nucleus to aid survival^17^, which might be a biological explanation of why in some cells and areas more MT-genes can be detected in sn-RNA-seq data in disease.

**Figure 2:**
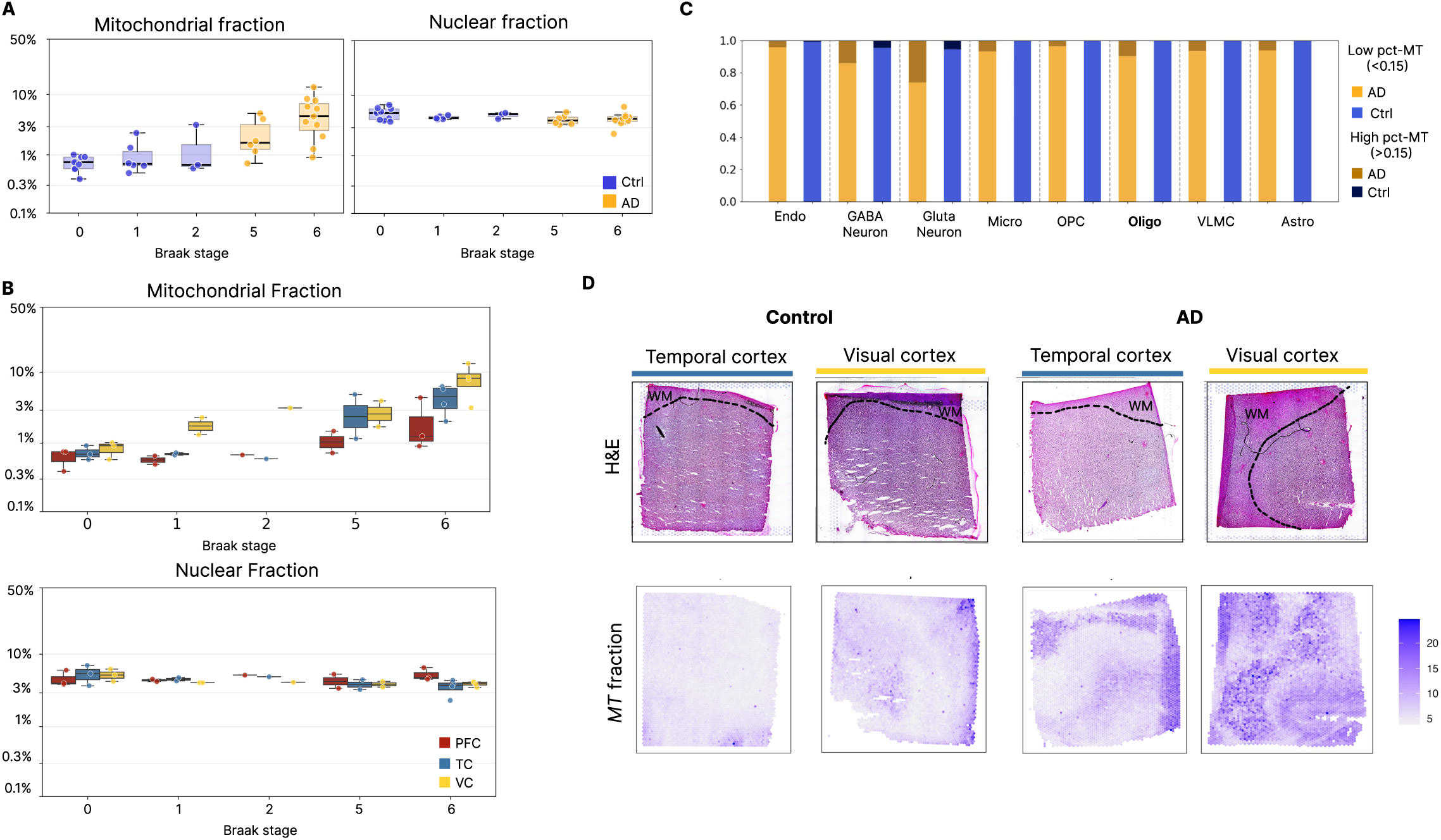
Mitochondrial gene expression varies across cortical areas and in disease. Pre-quality control analysis showing percentage of mitochondrial (*MT*) and nuclear fractions for all cells across **A** different Braak stages and **B** different cortical regions. **C** The percentage of low (<15%) and high (>15%) *MT* fraction shown per cell type and condition. **D** Visium analysis of temporal and visual cortices from control and AD brain samples. Upper panels show H&E staining with cortical layers and gray matter (GM) boundaries marked. Lower panels display spatial distribution of *MT* gene fractions across tissue sections.

Within all OL nuclei, we identified four unique OL states (Fig. 3A), of which none was exclusively associated with any singular sample variable, such as condition or cortical area (Fig. 3B), but we observed higher similarity between OL in TC and VC in comparison to PFC (Fig. 3C). This observation demonstrates global area-specific heterogeneity which might be related to OL function, in the case of promoting extracellular plasticity or circuit remodelling^18,19^. Interestingly, the PFC did not stand out in terms of composition of all cell types: for example GABAergic neuron composition mostly differed in the VC (Extended Data Fig 6). This finding highlights important structural differences between different cortical areas and exemplifies why analysing a single cortical brain region by snRNA-seq, as commonly practiced, and then extrapolating the results to other brain regions should be done cautiously.

**Figure 3:**
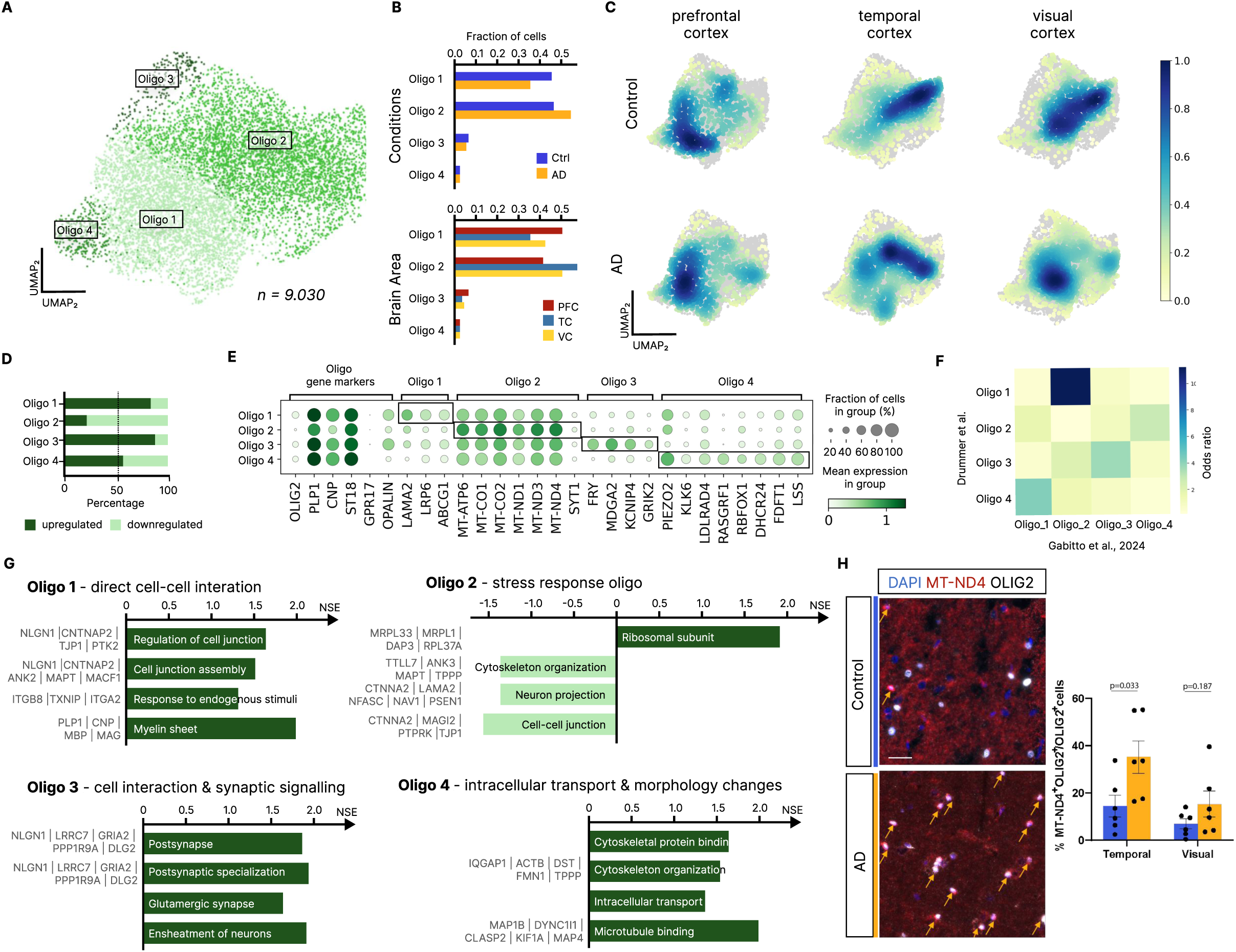
Oligodendrocyte subtypes vary across cortical areas and in disease. **A** UMAP plot of oligodendrocyte suptypes (n= 9030). **B** Bar plots depicting the fraction of cells per condition and area distributed across OL subtypes. **C** UMAP plots of OL with gaussian kernel density estimates over area and condition. **D** Bar plot depicting the percentage of up- and down-regulated differentially expressed genes (adjusted p-value < 0.05) across oligodendrocyte subtypes identified with DESeq2. **E** Dotplot showing oligodendrocyte subset markers based on DE analysis. **F** Heatmap showing the confusion matrix comparing OL annotations with a recently published reference dataset^16^. **G** Functional characterization of OL subclusters via pathway analysis, indicating normalized expression scores (p-value < 0.05) from each pathway; selected relevant genes of each pathway are displayed on the left. **H** Representative images and quantifications of MT-ND4+ OLIG2+ oligodendrocytes in the temporal and visual cortices of post-mortem human brains (n = 6 individuals per group). Scale bar = 25µm.

For each OL state, we identified up- and down-regulated genes (Fig. 3D) finding unique gene signatures per cluster (Fig. 3E): Oligo1 (4.557 nuclei, *LAMA2*, *LRP6*), Oligo2 (3.711 nuclei, *MT-ND4*, *MT-ND3*), Oligo3 (460 nuclei, *FRY*, *MDGA2*) and Oligo4 (199 nuclei, *PIEZO2*, *DHCR24*, *RBFOX1*). When compared to Gabitto et al., 2024^16^ that have the same OL subtype resolution, we observed a large agreement for cluster-specific upregulated genes (Fig. 3F). Gene Set Enrichment Analysis (GSEA)^20^ (Fig. 3G) of these clusters revealed functional differences, with Oligo1 being characterized by cell-cell interactions, Oligo3 by synaptic signalling and Oligo4 by morphological changes, cholesterol metabolism and intracellular transport. The latter RBFOX1-expressing Oligo4 cluster specifically mapped to the (immature) Oligo_1 from Gabitto et al. 2025 and was also reported in previous studies^21–23^. The genes related to high cholesterol metabolism point towards actively myelinating cells, the high abundance of *RBFOX1*, which is also highly expressed in neurons^24^ and in OL-neuron mixed lineage cells (Extended Data Fig. 2B), further pointing towards a neuromodulating function of Oligo4. Oligo2, on the other hand, which also showed an enrichment in AD (Fig. 3B), was predominantly characterized by the absence of cellular interactions, with a simultaneous upregulation of *MT* genes, as highlighted in previous research^21,22^. The aforementioned *MT*-gene upregulation is associated with disease and might result in the loss of important cellular functions, for instance, a loss of OL-axonal coupling or myelin maintenance. MT-ND4 in particular has recently been suggested as a promising biomarker of AD^25^. IHC validations in a different patient cohort corroborated a higher abundance of MT-ND4+ OL in the TC of AD subjects as compared to controls (Fig. 3H). In contrast, other prominent disease-associated OL markers from the literature^26–29^, such as *SERPINA3*, could not be observed (Extended Data Fig. 7). While this could be due to technical reasons (e.g. sequencing depth or single nucleus vs single cell analysis)^30^, we hypothesize that these markers are species-specific, as *SERPINA3* has predominately been reported in mouse models and not consistently in humans^7,22^.

In conclusion, our work reveals functional differences between OL states, which in AD show a loss of essential functions such as myelin maintenance, axonal support, and metabolic processes. We identify mitochondrial activation as a general hallmark of disease progression, demonstrating that up-regulation of mitochondrial genes is not merely a technical artifact but reflects underlying pathology. Additionally, we show that the loss of characteristic cell signatures in AD suggests widespread changes in cellular functions across multiple cell types. Together, these findings advance our understanding of the cellular mechanisms driving AD pathogenesis and lay the groundwork for developing targeted therapies to restore oligodendrocyte health and function.

## Methods

### Human subjects

Post-mortem tissue for this project was obtained from The Edinburgh brain bank, the Newcastle brain bank and the Munich brain bank with ethical approval from the Ethics Committee at the LMU Munich (vote 22-1060).

| Donor ID | Condition | Sex | Age | Tissue Source |
| --- | --- | --- | --- | --- |
| Frozen Tissue samples for snRNAseq and Visium analysis |  |  |  |  |
| Ctrl3 | Control | male | 70 | Neurobiobank Munich |
| Ctrl4 | Control | male | 76 | Neurobiobank Munich |
| 20130894 | Control | male | 80 | Newcastle Brain Resource |
| 20110891 | Control | male | 73 | Newcastle Brain Resource |
| 20100742 | Control | male | 67 | Newcastle Brain Resource |
| SD030/18 | Control | male | 63 | Edinburgh Brain Bank |
| SD006/14 | Control | male | 60 | Edinburgh Brain Bank |
| 20185928 | Alzheimer's Disease | male | 69 | Newcastle Brain Resource |
| 20174945 | Alzheimer's Disease | male | 69 | Newcastle Brain Resource |
| AD3 | Alzheimer's Disease | male | 70 | Neurobiobank Munich |
| AD5 | Alzheimer's Disease | male | 74 | Neurobiobank Munich |
| AD6 | Alzheimer's Disease | male | 77 | Neurobiobank Munich |
| AD7 | Alzheimer's Disease | male | 75 | Neurobiobank Munich |

### snRNA-seq pipeline

#### Nuclei sampling and library preparation

Fresh-Frozen tissue blocks were cut into 20µm sections and Luxol-Fast-Blue stainings were performed to determine tissue architecture. Gray Matter regions from ~10-20 sections per sample were collected using a razor blade. Nuclei isolation was performed with a nuclei isolation kit (Nuclei EZ Prep; NUC101; Sigma Aldrich) according to the manufacturers instructions and nuclei underwent FACS sorting for NeuN (FCMAB317PE; Merck) and DAPI. To enrich our samples for GM cells, we collected 10.000 DAPI+NeuN+ nuclei and 10.000 DAPI+NeuN− negative nuclei per sample to precede with the library preparation.

#### Library preparation and Sequencing

Samples were randomly distributed to 10X Genomics Chromium single cell 3’ chips. LIbraries were prepared with the 10X Genomics v3 beads, Gem kits and library kits, according to the manufacturer’s instructions. Library quality and final concentrations were determined using a Bioanalyzer.

For sequencing, libraries were again randomized and 9-10 libraries were combined at equimolar concentrations and adjusted to 2nM for sequencing on a NovaSeq S4 Flowcell with a targeted sequencing depth of 50K reads per nucleus at the Competence Centre for Genomic Analysis in Kiel. To avoid sequencing batch effects, each pool was sequenced twice on different lanes and the results were combined.

#### snRNA-seq preprocessing, QC and annotation

We followed the best practices single-cell analysis suggestions from Heumos et al., 2023^12^ and their updated online resource: https://www.sc-best-practices.org/preamble.html.

#### snRNA-seq data preprocessing

Gene counts were obtained by aligning reads to the GRCh38 (hg38) Genome Ensembl 105 using Cell Ranger software (v.9.0.0) (10x Genomics). The filtered cell ranger matrix contains 42.540 cells and 78.932 genes (Supp. Table 1). First, we filter low quality cells given cell clusters that are associated with low number of detected genes, low count depth or high fraction of mitochondrial counts using Scanpy^31^. Additionally, we consider *MALAT1* expression of the cells which is related to the nuclear fraction of cells captured^32^. During this preprocessing step we mainly remove cells from the three samples AD3 - GR, AD5 - AS and Ctrl6 - GR that are associated with low number of cell counts and genes according to cellranger (Supp. Table 1) and distribution plots (Suppl. Fig 1).

From initial observations there are expression and count differences between condition and brain area samples that we suspect to be biological rather than technical (difference in mean but rather stable standard deviation). Therefore, we apply sub-group specific thresholds per condition and brain area for a biology inspired data-driven quality control^14^ (Suppl. Tab 2). To account for ambient RNA we corrected gene counts using SoupX^33^, removing 44.117 genes and we removed 898 cells from high count doublet clusters (>0.9) detected by scublet^34^. Another cluster with higher than average doublet count was kept, because it contains OPCs that have a higher transitional profile which are possibly mistaken as doublets and the lower amount of doublets expected in single-nuclei data.

#### Normalization and dimensionality reduction

Functions in the Scanpy^31^ (v1.10.4) package were used for the following analyses. First we selected highly variable genes (2k) and then normalised the counts to the median of total counts for observations (cells) before normalization. As suggested by a recent benchmark^35^, we applied a shifted logarithm transformation 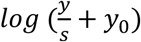 on the raw counts *y* with a size factor *s* and pseudo-count *y*_0_. After scaling the data, we performed Principal Component Analysis (PCA) with 50 PCs followed by Uniform manifold approximation and projection (UMAP) based on their 15 neighborhood embeddings.

#### Batch analysis and integration

For integration, we first determined which covariates cause batch effects in the dataset. We included technical (e.i. brain bank source, donor) and biological covariates (e.i. Braak stage, cortex area) in the analysis. To quantify the contribution of covariates we performed Principal Component Regression (PCR) using scAtlasTB^36^ (https://github.com/HCA-integration/scAtlasTb/). We considered four covariates and computed the variance explained (PRC) by each covariate across the top principal components of the gene expression matrix, comparing observed values against a null distribution generated by permutation (N=30). Significant covariates were identified based on z-scores and empirical p-values. The donor identity accounted for the largest variance (pcr = 0.045, z = 4.48, n = 12, p < 0.01), followed by brainbank source (pcr = 0.013, z = 3.50, n = 3, p < 0.01), Braak stage (pcr = 0.010, z = 3.09, n = 5, p < 0.01), cortex area (pcr = 0.010, z = 2.52, n = 3, p < 0.05), and condition (pcr = 0.009, z = 2.87, n = 2, p < 0.05).

To remove the technical batch effects we integrated on donor and brain bank sources using scVI^37^, scANVI^38^. We evaluate the data integration with metrics from scIB^39,40^ to compare biological and technical batch removal. Finally, the scVI integrated data on donors was chosen as the most meaningful representation.

#### Clustering and cell type annotation

Clustree^41^ (v.0.4.1) was used to select the most stable clustering resolution (res=1.0) for the overall. Using sc.tl.leiden with resolution 1.0 we obtained 28 clusters on the full dataset. Cluster 26 and 27 were removed due to low cell counts (8 and 39 cells respectively) (see Supp. Table 2). To identify major cell types, we assigned clusters to cell types according to cell-type specific markers from literature ^6,7,21,42^. Next we saved the raw count data in .h5ad format and the deep generative mapping algorithm from MapMyCells (RRID: SCR_024672) was used to align the nuclei with the 10x human MTG reference (CCN20230505). The mapping results were projected onto the leiden clustering and a cluster was assigned to a cell type if the mapped fraction of one cell type was higher than 0.75 (Suppl. Table 4). Most mapped annotations agree with the manual annotations. A subset of 2.670 cells were not able to be associated with a specific cell type cluster, indicating mixed-lineage clusters (Suppl. Figure 3). According to the cell type mapping, this subset mainly seems to consist of Glutamatergic Neurons, Oligodendrocytes, Microglia and Astrocytes.

The final data cohort consists of a total of 26.598 nuclei across 6 controls (15.270 cells) and 6 AD patients (11.328 cells) with 34.802 genes, with a median of 10.354 counts and 3.086 features per nucleus.

#### Differential gene expression analysis

Differential gene expression analysis was performed using the DESeq2^43^ package (v.1.48.1) in R (v.4.5.1). Gene expression count matrices and associated metadata were prepared from single-cell RNA sequencing data. To avoid issues with zero counts, a pseudocount of 1 was added to all expression values and low expression genes were filtered if they had at less than 10 counts in a minimum of 3 samples. We performed i) one-vs-all and ii) pairwise comparisons. For the one-vs-all comparison, each experimental condition was compared against all other conditions combined while the pairwise comparison compared conditions (AD, Ctrl) in the same biological context (e.i. cortical areas).

Results were summarized to report the total number of genes tested, significantly upregulated genes (positive log2 fold change, FDR < 0.05), and significantly downregulated genes (negative log2 fold change, FDR < 0.05) for each comparison. Furthermore, we visualize the results with volcano plots showing the change in expression on the x-axis (log-fold-change) and statistical significance on the y-axis (FDR-corrected p-values). We color code the genes that have a FDR-corrected p-value under 0.01 and log-fold-change over 1.5.

#### Gene Set Enrichment Analysis

Functional annotation of gene sets was performed with Fast Gene Set Enrichment Analysis (fgsea)^20^ on each oligodendrocyte cluster using the differentially expressed gene lists. The gene pathways from Molecular Signatures Database (MSigDB)^44^ were used specifically H, C2 and C5, which is the gene ontology collection. For each oligodendrocyte cluster we pre-filtered pathways with a p-value < 0.05 and then selected 4 representative pathways for each cluster with associated genes.

#### Compositional Analysis of cell types

We ran scCoda^45^ for the compositional analysis using the pertpy framework (v1.0.0)^46^ to compare the relative abundance of major cell types. We ran the analysis across four different covariate combinations: 1) condition across all brain areas and 2) within brain areas, 2) brain region and 3) combined condition and brain area. The recommended false discovery rate (FDR) value from pertpy for the scCoda analysis is 0.2. To receive credible effects for condition (AD vs Ctrl) analysis (1,2) we needed to increase the FDR to 0.5. For the other covariates that involve cortex area differences a FDR of 0.2 was sufficient. Because there is no consensus on which supertypes would be affected by AD, we ran the compositional analysis with each cell type set as the unchanged reference population, as recommended by the authors of scCODA. The final results were averaged across reference cell types (Supp. Table 3).

#### Canonical marker gene expression differences

For canonical marker genes to accurately identify cell types their expression must be high in the cell type clusters. As cell type annotation mapping was performed before integration some clusters contain an unbalanced amount of Ctrl vs AD cells. We observed a general trend that clusters with a higher Ctrl fraction could be more accurately mapped. One potential reason could be a decrease of canonical marker gene expression with an increase of disease related genes that are also considered canonical marker genes (e.i. CLU - astrocyte marker but also a myelination gene that is known to decrease during AD progression). We compare canonical gene marker expression sample wise between conditions (Fig. 1G).

#### Mitochondrial and nuclear gene fraction processing

Gene expression fractions were calculated at the sample level by aggregating specific gene sets and total gene counts across all cells within each sample. To improve visualization of differences in low fraction ranges where most samples typically fall and numerical stability, we applied a logit transformation following the formula: logit_fraction = logit((target_genes + 1)/(total_genes + 2)), where target_genes represents the sum of counts for the gene set of interest and total_genes represents the sum of total gene counts per sample, and added pseudocounts.

Data were visualized using grouped boxplots with individual sample points overlaid, stratified by experimental conditions. The logit-transformed y-axis was displayed with percentage labels (0.1%, 0.3%, 1%, 3%, 10%, 50%) to facilitate interpretation while maintaining the improved resolution in the low percentage range.

For mitochondrial fraction analysis, mitochondrial genes comprised 37 mitochondrial-encoded genes (*MT-TF, MT-RNR1, MT-TV, MT-RNR2, MT-TL1, MT-ND1, MT-TI, MT-TQ, MT-TM, MT-ND2, MT-TW, MT-TA, MT-TN, MT-TC, MT-TY, MT-CO1, MT-TS1, MT-TD, MT-CO2, MT-TK, MT-ATP8, MT-ATP6, MT-CO3, MT-TG, MT-ND3, MT-TR, MT-ND4L, MT-ND4, MT-TH, MT-TS2, MT-TL2, MT-ND5, MT-ND6, MT-TE, MT-CYB, MT-TT, MT-TP*). For nuclear fraction analysis, nuclear markers included 30 nuclear-encoded genes (*MALAT1, NEAT1, FTX, FOXP1, RBMS3, ZBTB20, LRMDA, PBX1, ITPR2, AUTS2, TTC28, BNC2, EXOC4, RORA, PRKG1, ARID1B, PARD3B, GPHN, N4BP2L2, PKHD1L1, EXOC6B, FBXL7, MED13L, TBC1D5, IMMP2L, SYNE1, RERE, MBD5, EXT1, WWOX*).

#### Oligodendrocyte subclusters comparison with Gabitto et al., 2024

To compare the oligodendrocyte subclusters with the oligodendrocyte subclusters from Gabitto et al., 2024^16^ we used the up- and downregulated DE genes identified per cluster. The DE genes in Gabitto et al., 2024 were calculated with Nebula^47^ while we used DESeq2.

To systematically assess the overlap between DE gene signatures, we performed enrichment analysis using Fisher’s exact test. First, we separated genes into upregulated and downregulated sets based on their log fold change values. We defined the analysis universe as the intersection of all genes detected in both datasets to ensure appropriate statistical comparison. For each direction combination (up-up, up-down, down-up, down-down), we calculated the overlap between gene sets and performed one-sided Fisher’s exact tests and calculate the odds ratios to quantify the strength of association between gene sets and applied false discovery rate (FDR) correction using the Benjamini-Hochberg procedure to account for multiple testing across the four directional comparisons.

### Visium analysis

#### Visium pipeline and sequencing

For the visium pipeline, 4 of the best samples from the snRNA-seq experiment were chosen and small blocks were further processed for the 10x Genomics Visium Spatial Gene Expression pipeline and the protocol was performed according to the manufacturer’s instructions. Libraries were assessed by a bioanalyzer and sequenced with a NextSeq 2000 P3 Flowcell with a sequencing depth of 400K reads per sample.

#### Quality control

For spatial validation of the condition and brain area specific effects we obtained data from one Visium slide with 4 images from one control and one AD patient for each temporal and visual cortex respectively. Gene counts were obtained by aligning reads to the GRCh38 (hg38) Genome Ensembl 105 using Space Ranger software (v.2.1.1) (10x Genomics). We performed basic quality control (removed zero count genes) followed by a locally aware quality control using SpotSweeper^48^ to account for region specific trends per sample independently. For further statistical testing more replicates are needed, hence we are only using it as support for our other findings.

The MT fraction was calculated using a subset of 13 MT-genes in Visium data from the snRNA-seq data: *MT-ATP6, MT-ATP8, MT-CO1, MT-CO2, MT-CO3, MT-CYB, MT-ND1, MT-ND2, MT-ND3, MT-ND4, MT-ND4L, MT-ND5, MT-ND6*.

#### Immunohistochemistry oh post-mortem brain samples

4µm formalin-fixed paraffin-embedded tissue sections were first deparaffinized in 100% RotiHistol twice for 5 minutes and rehydrated by incubating in decreasing concentrations of ethanol (100% twice, 96%, 80%, 70%) for 1 minute each. Sections were then briefly washed with water and boiled for 10 minutes in pre-heated antigen unmasking solution buffer (Vector Laboratories, H-3300, 1:100). After a brief wash, autofluorescence was quenched with an autofluorescence eliminator reagent (EMD Millipore, 2160) for 1 minute followed by a 10-minute incubation with freshly made 3% H2O2, a wash in Tris-buffered saline + 0.5% Triton (TBS-T) and a 1-hour incubation in blocking solution (TBS-T, 10% heat-inactivated horse serum). Sections were incubated in primary antibody diluted in blocking solution at 4°C overnight in a humid chamber. Primary antibodies used for immunohistochemistry were: anti-OLIG2 (rabbit, Sigma-Aldrich, HPA003254, 1:100), anti-Olig2 (goat, R&D Systems, AF2418, 1:200), anti-CNP (mouse, Sigma-Aldrich, AMAB91072, 1:1000), anti-MT-ND4 (rabbit, Invitrogen, PA5-116791, 1:200), anti-AACT (SERPINA3) (rabbit, Abcam, AB205198, 1:1000), anti-CD31 (goat, R&D Systems, AF3628-SP, 1:100). The next day the sections were washed in TBS-T and incubated for 2 hours with the matching secondary peroxidase-conjugated antibody. Secondary antibodies that were used: ImmPRESS IgG Polymer Kits, Peroxidase, Vector Laboratories (horse anti-goat, MP-7405; horse anti-rabbit, MP-7401; horse anti-mouse, MP-7402). After washing the sections in TBS-T, they were incubated with fluorophore diluted 1:500 in borate buffer solution (100 mM borate buffer, 0.003% H2O2) for 10 minutes. The fluorophores used were TSA Vivid Fluorophore 520, 570 and 650 (Bio-Techne, 7523/1, 7526/1, 7527/1). Multiplexed stainings were done sequentially, performing the incubation of each primary antibody separately overnight, after repeating the antigen retrieval step with the boiling time reduced to 3 minutes. Nuclei were counterstained with DAPI diluted 1:1000 in PBS for 3 minutes and slides were washed in PBS and mounted with Fluoromount-G (0100-01, Southern Biotech).

Images of the entire human brain sections were acquired with the Pannoramic Midi II Slide Scanner (3D HISTEC) with the 20x objective. Scans were converted into tiff-files using the NGFF-Converter (Glencoe Software) and for each patient, approximately 10 random images of similar size were selected per each cortical layer set and cells were quantified manually or semi-quantitatively with the cell-counter plugin.

#### Statistics

No statistical methods were used to predetermine sample sizes, but our sample size for the snRNA-seq and the validation experiments was comparable to previously published studies. For the snRNA-seq analyses Python- and R-based open source packages were used and the statistical analysis is described in the dedicated methods section. For the immunohistochemical validations, graphical visualizations and statistical analysis were performed using GraphPad Prism software (10.3.1). Differences between two groups were determined using a student’s t-test. For statistically significant comparisons (p<0.05) exact p-values are displayed in the figures, all other comparisons did not show a significant p-value.

## Data availability

We obtained preprocessed reads from one publicly available dataset that performed single-nucleus RNA-seq in AD patients (Gabitto et. al, 2024)^16^. This dataset can be accessed through AWS (https://sea-ad-single-cell-profiling.s3.amazonaws.com/index.html#MTG/RNAseq/Supplementary%20Information/Nebula%20Results/.)

Our own generated (snRNA-seq and Visium) data will be made available upon acceptance of the manuscript.

## Code availability

The collection of scripts used to annotate the snRNA-seq & Visium data and publicly available datasets, perform all the analyses and create each figure are available at the Jaekel lab github page: https://github.com/Jaekellab/human_AD_snRNAseq/tree/main.

## Acknowledgments

We want to thank the following people for their contribution to the manuscript: Felix Fischer for setting up the analysis and the notebook structure for the bioinformatic analysis. Vladimir Shitov and Daniel Strobl for the guidance on the cellranger output. Michaela Müller for the help with integration and access to the scAtlasTab pipeline. Will Macnair and Sergio Marco-Salas for the helpful feedback on the figures and the manuscript. All the developers of the tools that have been used and the sc-best-practices consortium for the tutorials and environment files. Finally, we thank all the member from the group of FT and SJ for their insightful discussions, feedback, and support.

## Author contributions

S.J. designed the study and together with F.J.T. supervised the project. C.H. performed snRNA and visium experiments and performed the library preparations with the help of E.B. F.D. wrote the code and performed bioinformatic data analysis under the supervision of F.J.T. and L.K. S.J. and F.D. interpreted the results from the data analysis; N.D. and S.J. performed experimental validations. F.D designed and created the figures and together with S.J. wrote the manuscript. All authors read, corrected and approved the manuscript.

## Competing interests

F.J.T. consults for Immunai, CytoReason, BioTuring, Genbio and Valinor Industries, and has ownership interest in RN.AI Therapeutics, Dermagnostix, and Cellarity. S.J. has a financially supported research collaboration with Ono Pharmaceuticals.

## Funding

S.J. and her research were funded by the DFG Emmy-Noether Programme (JA 2891/2-1) and the Hertie Network of Excellence in Clinical Neuroscience. F.J.T received the following funding: DeepCell: Funded by the European Union (ERC, DeepCell - 101054957), HCA Data Ecosystems: This work was supported by the Chan Zuckerberg Initiative Foundation (CZIF; grant CZIF2022-007488 (Human Cell Atlas Data Ecosystem)), MCML: This work was supported by the German Federal Ministry of Research, Technology and Space (BMFTR) under grant no. 01IS18053A. Leibnizpreis: This work was supported by the DFG Leibniz Prize awarded to F.J.T.

## Figure legends

**Extended Data Figure 1: General quality control metavariables to determine outliers**.

**A-C** Violin plots showing the number of identified counts and genes (**A**), mitochondrial fractions (**B**) and *MALAT1* expression (**C**) across all samples. Abbreviations: AS - area striata (part of the visual cortex), GR - Gyrus Rectus (part of the prefrontal cortex, TG - Temporal Gyrus (part of the temporal cortex). Samples highlighted by a red rectangle were removed for further analysis. **D** UMAP plots of leiden clusters (res=1.0), log1p_counts, *MALAT1*-expression and *MT-*fraction over the entire dataset for quality control.

**Extended Data Figure 2: Mixed lineage cluster cells express oligodendrocyte, astrocyte and neuronal marker genes.**

**A** Unannotated cells (n=2670) highlighted in the UMAP plot of the complete, quality controlled data. **B** Dotplot of canonical marker genes in these unannotated cells demonstrating the expression of mixed-lineage markers, emphasizing astrocyte (*SLC1A2*), oligodendrocyte (*MBP*, *PLP1*) and Inhibitory neurons (*SYT1*) markers. Additionally, previously reported mixed-lineage markers^3^ (*RBFOX1*, *PDE4B*) are expressed. **C** The integrated mixed-lineage cluster UMAPs colored by condition and cortical area show higher separation between conditions than areas. **D** UMAP overlay of MT and nuclear fractions in unannotated cells. **E** Examples of immunohistochemical stainings of individual cells in AD brains showing co-labelling of the EC marker CD31 and the mature OL marker CNP (orange arrows). CNP+ OL that are CD31- are indicated by green arrows. Scale bars: 10µm.

**Extended Data Figure 3: Canonical marker gene expression is changed in individual cell types in lzheimer’s disease.**

**A** Quantifications of OLIG2 and CNP+ cells in the TC and VC in Control and AD tissue. **B** Representative images and quantifications of CD31+ EC in the temporal cortex in control and AD. Scale bar = 50µm. **C** Comparison of the sample-wise mean expression of canonical marker genes for each cell type between control and AD.

**Extended Data Figure 4: *MT* - and nuclear fractions vary across different conditions.**

**A** Percentage of *MT* fraction and *MALAT1* expression for all cells across condition and cortical area in this dataset. **B** Quality control filtering matrix shows the number of cells removed by different thresholds, based on number of genes detected, UMI counts, mitochondrial fraction, and *MALAT1* expression, across conditions and cortical areas. **C** Percentage of *MT* and nuclear fractions for all cells across Braak stages and regions from Gabitto et al., 2024.

**Extended Data Figure 5: Quality control results are more confounded with regional differences in AD slides than control slides.**

Quality control analysis with SpotSweeper^48^ providing results for global and local outliers. Global outliers in slides from a control (**A**) and AD patient (**B**) seem to follow coherent patterns rather than random allocations suggesting that high thresholds pick up on regional differences. For each control and AD subject two slides from temporal and visual cortices are shown with log2 total and detected counts. From the results we can show that no slide shows artifacts of 1) dryspots - not visible in histological images, identified by low library sizes and no difference in mitochondrial ratio or 2) hangnail artefacts - library sizes and no difference in mitochondrial ratio.

**Extended Data Figure 6: Regional density distributions per cell type mirror Alzheimer’s disease progression gradient.**

UMAPs showing the kernel dentistry estimation (scanpy.tl.embedding_density) in all broad cell types (besides OL) across the temporal, prefrontal, and visual cortex. Regional gradients reflect decreasing AD pathology from lowest (visual cortex) to highest pathology (temporal cortex), based on pathological assumptions thereby providing a spatial snapshot of cellular changes along disease progression.

**Extended Data Figure 7: Common disease-associated oligodendrocyte markers are not expressed.**

**A** Dotplot showing the log-scales mean expression of common disease-associated OL (DAO) genes taken from the literature and divided into four groups: 1) transcription factors (TFs), 2) reactive markers, 3) immune-response genes and 4) genes associated with cholesterol metabolism. While TF expression might be lower in snRNA-seq vs scRNA-seq, other frequently cited DAO genes, such as *SERPINA3*, also do not show high expression. **B** Representative images and quantifications of immunohistochemical stainings of SERPINA3+OLIG2+ OL in human brains across cortical layers in the temporal cortex demonstrating a low abundance of SERPINA3+ OL and no specific enrichment in AD. Scale bars: 50 µm. **C-E** Analysis of *SERPINA3* expression in the temporal cortex; snRNA-seq data from Gabitto et al., 2025. UMAP of all cell types (OL solid circle, astrocytes dashed circle) (**C**). UMAP overlay of *SERPINA3* expression across celltypes, with the highest enrichment observed in astrocytes rather than oligodendrocytes (**D**). Dotplot showing commonly reported DAO genes in OL, supporting the observation that *SERPINA3* is not highly expressed in disease OL (**E**). **F-H** Parallel analysis of SERPINA3 expression in dorsolateral prefrontal cortex (DLPFC); snRNA-seq data from Gabitto et al., 2025. UMAP of all cell types (OL solid circle, astrocytes dashed circle) (**F**). UMAP overlay of *SERPINA3* expression across cell types, with the highest enrichment observed in astrocytes rather than oligodendrocytes, consistent with temporal cortex findings (**G**). Dotplot showing oligodendrocyte-specific marker gene expression in DLPFC, confirming the absence of SERPINA3 expression in oligodendrocytes across different cortical regions (**H**).

## Notes

https://github.com/Jaekellab/human_AD_snRNAseq

### Competing Interest Statement

The authors have declared no competing interest.

### Summary of Updates

Author order adjusted: Sarah Jaekel as last author

