## Extended Figures and Tables for "Brain Region-Specific Oligodendrocyte States Highlight Mitochondrial Gene Upregulation and Loss of Canonical Identity Signatures in Alzheimer’s Disease"

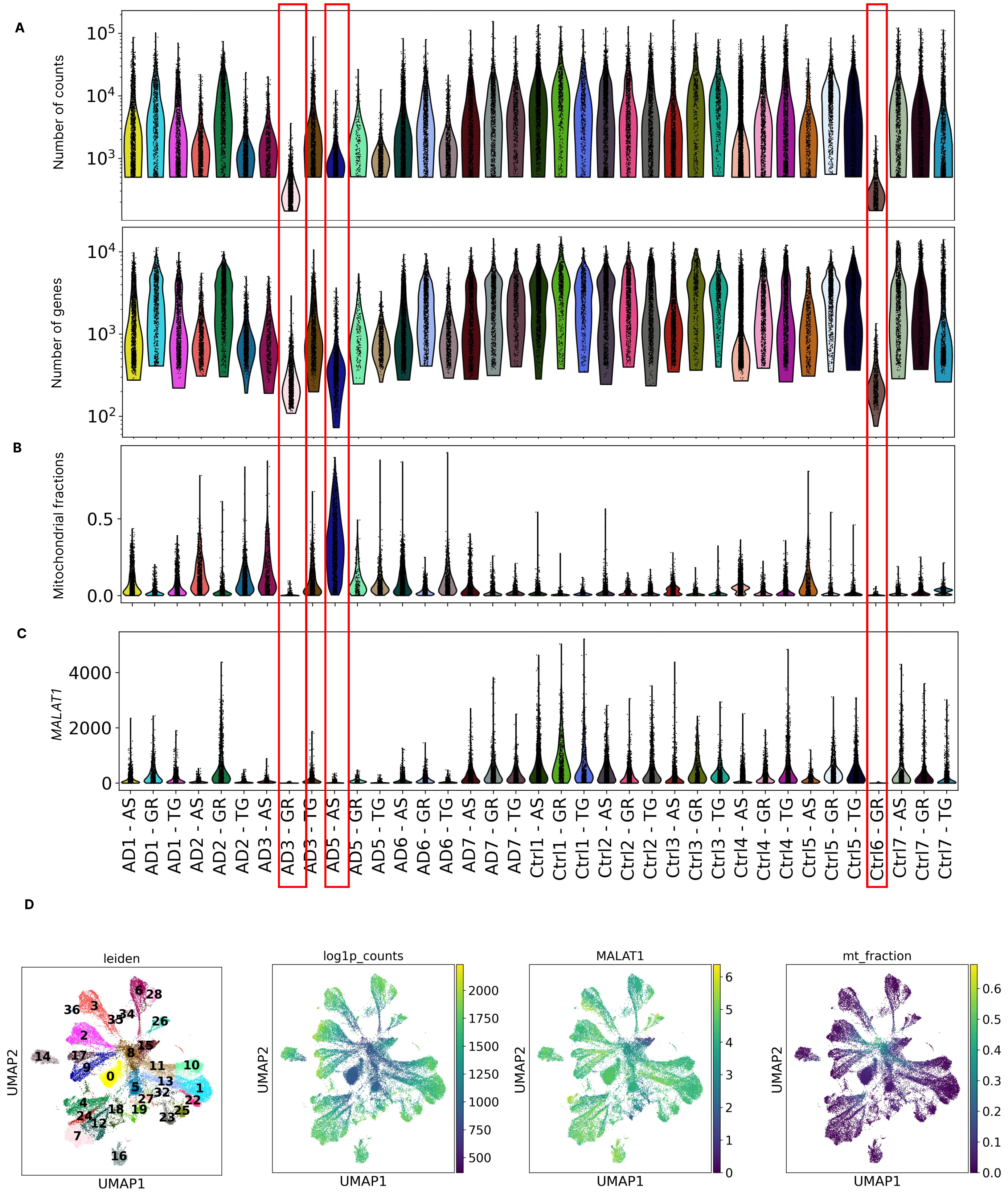


**Extended Data Figure 1: General quality control metavariables to determine outliers.**

**A-C** Violin plots showing the number of identified counts and genes (**A**), mitochondrial fractions (**B**) and MALAT1 expression (**C**) across all samples. Abbreviations: AS - area striata (part of the visual cortex), GR - Gyrus Rectus (part of the prefrontal cortex, TG - Temporal Gyrus (part of the temporal cortex). Samples highlighted by a red rectangle were removed for further analysis. **D** UMAP plots of leiden clusters (res=1.0), log1p_counts, MALAT1-expression and MT-fraction over the entire dataset for quality control.

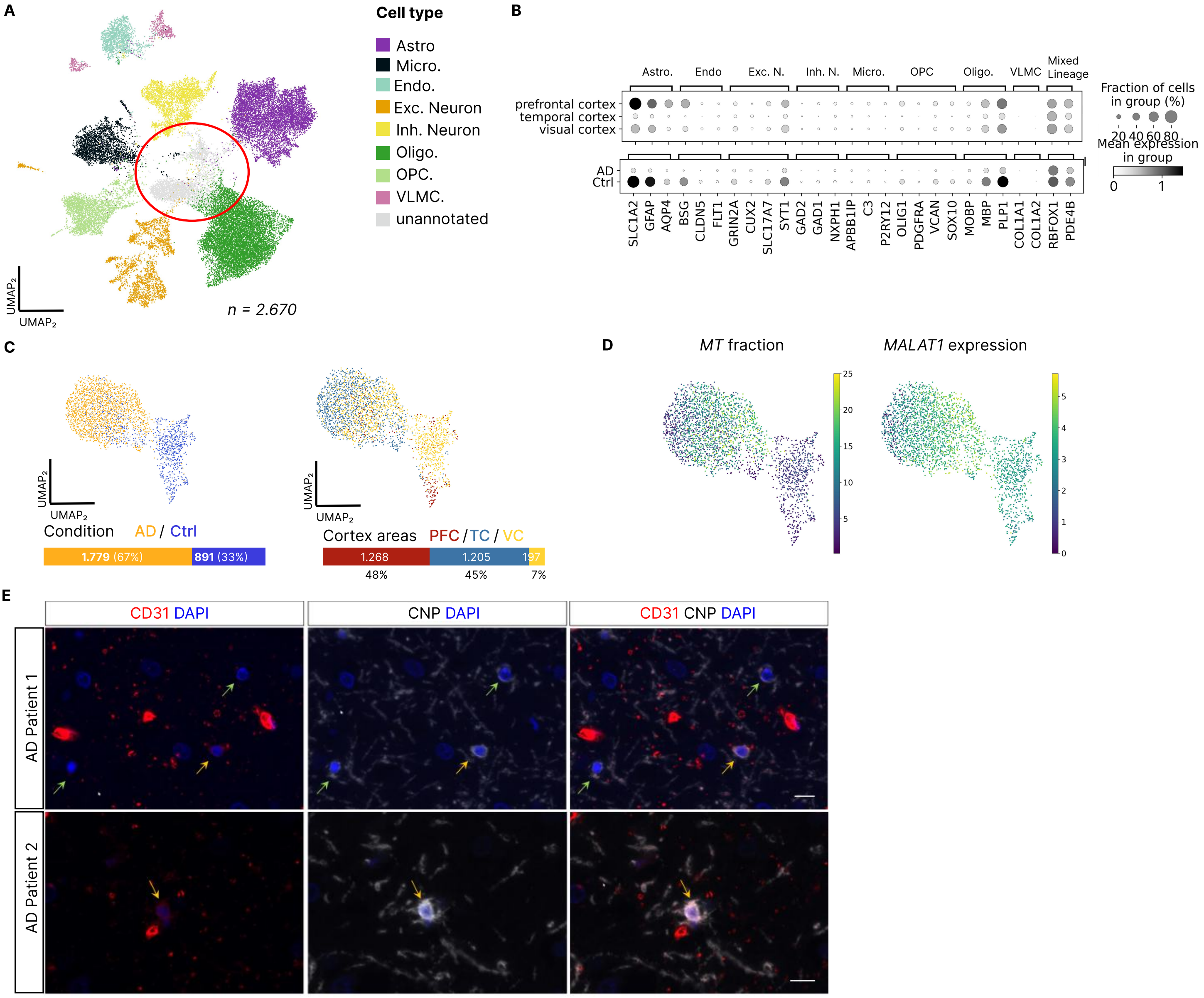


**Extended Data Figure 2: Mixed lineage cluster cells express oligodendrocyte, astrocyte and neuronal marker genes.**

**A** Unannotated cells (n=2670) highlighted in the UMAP plot of the complete, quality controlled data. **B** Dotplot of canonical marker genes in these unannotated cells demonstrating the expression of mixed-lineage markers, emphasizing astrocyte (SLC1A2), oligodendrocyte (MBP, PLP1) and Inhibitory neurons (SYT1) markers. Additionally, previously reported mixed-lineage markers3 (RBFOX1, PDE4B) are  expressed. **C** The integrated mixed-lineage cluster UMAPs colored by condition and cortical area show higher separation between conditions than areas. **D** UMAP overlay of MT and nuclear fractions in unannotated cells. **E** Examples of immunohistochemical stainings of individual cells in AD brains showing co-labelling of the EC marker CD31 and the mature OL marker CNP (orange arrows). CNP+ OL that are CD31- are indicated by green arrows. Scale bars: 10µm.

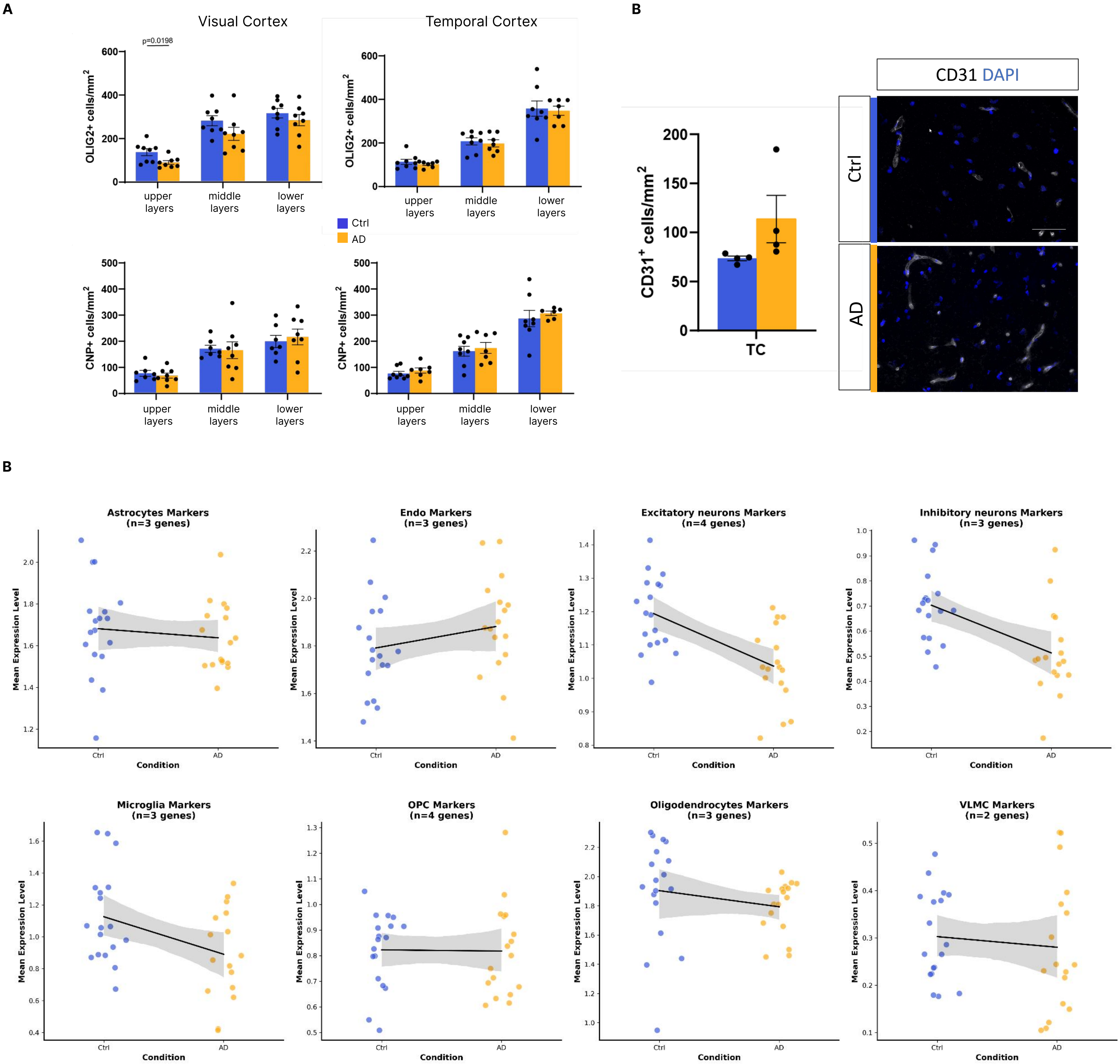


**Extended Data Figure 3: Canonical marker gene expression is changed in individual cell types in lzheimer's disease.**

**A** Quantifications of OLIG2 and CNP+ cells in the TC and VC in Control and AD tissue. **B** Representative images and quantifications of CD31+ EC in the temporal cortex in control and AD. Scale bar = 50µm. **C** Comparison of the sample-wise mean expression of canonical marker genes for each cell type between control and AD.

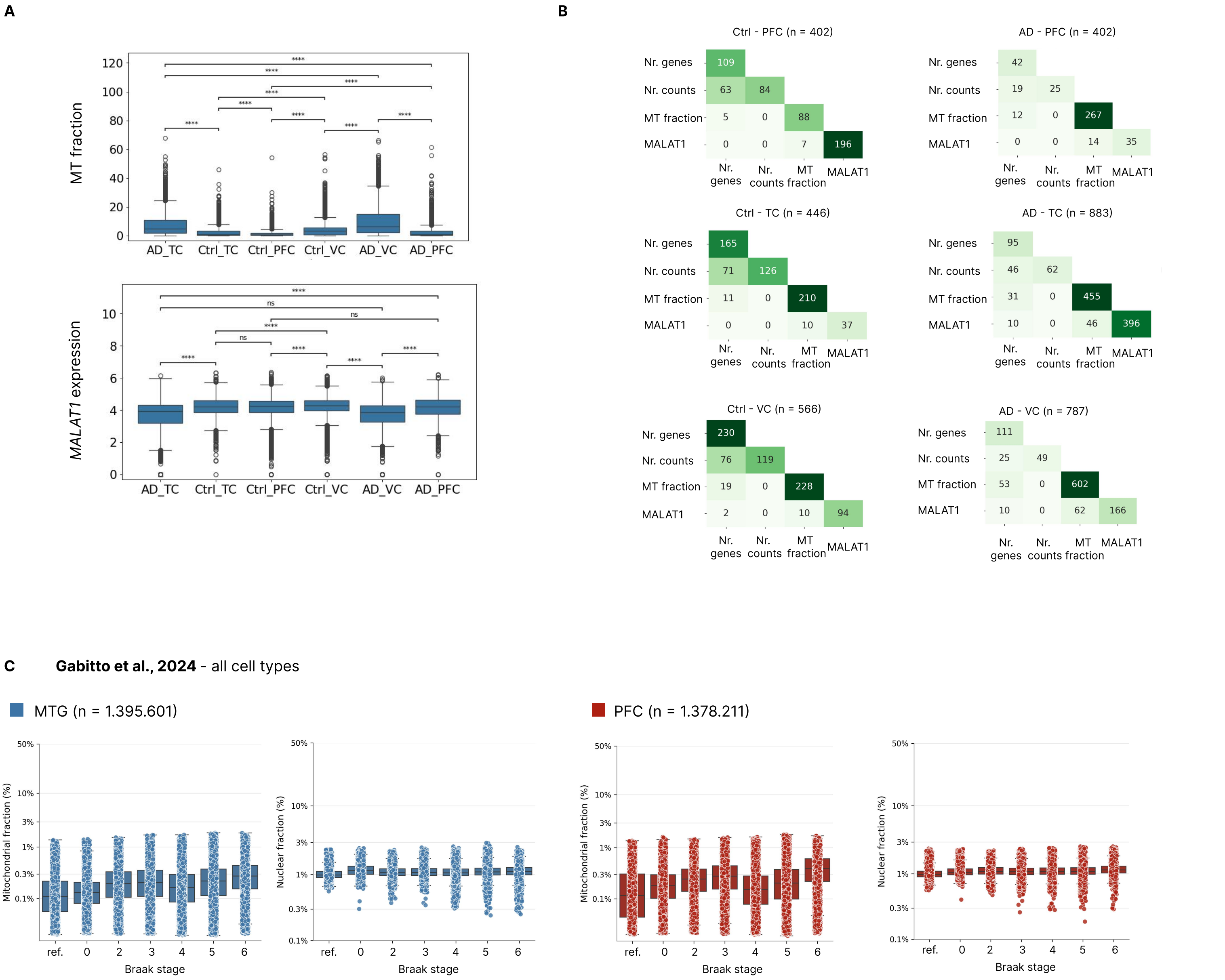


**Extended Data Figure 4: MT - and nuclear fractions vary across different conditions.**

**A** Percentage of MT fraction and MALAT1 expression for all cells across condition and cortical area in this dataset. **B** Quality control filtering matrix shows the number of cells removed by different thresholds, based on number of genes detected, UMI counts, mitochondrial fraction, and MALAT1 expression, across conditions and cortical areas. **C** Percentage of MT and nuclear fractions for all cells across Braak stages and regions from Gabitto et al., 2024.

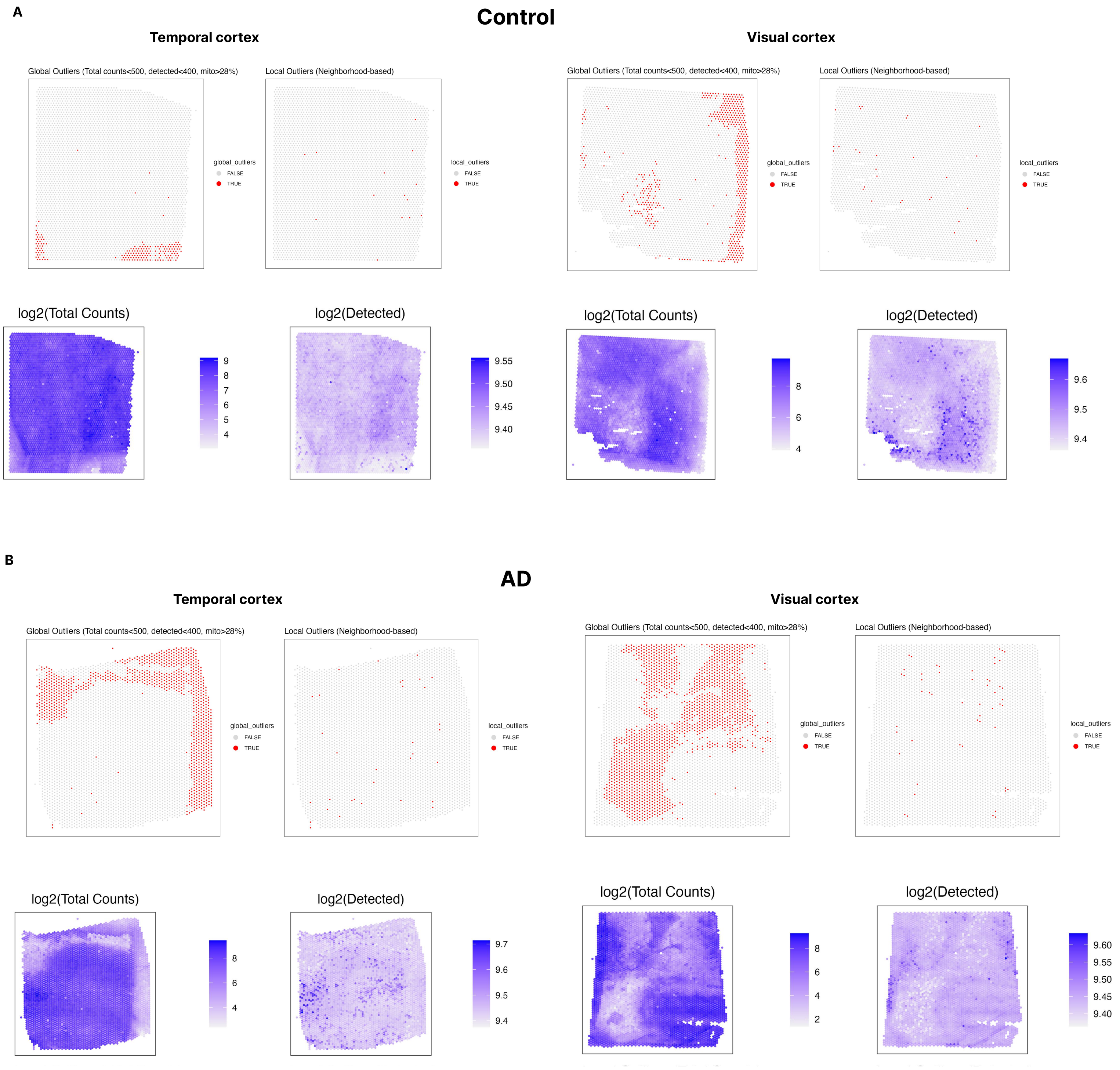


**Extended Data Figure 5: Quality control results are more confounded with regional differences in AD slides than control slides.**

Quality control analysis with SpotSweeper48 providing results for global and local outliers. Global outliers in slides from a control (**A**) and AD patient (**B**) seem to follow coherent patterns rather than random allocations suggesting that high thresholds pick up on regional differences. For each control and AD subject two slides from temporal and visual cortices are shown with log2 total and detected counts. From the results we can show that no slide shows artifacts of 1) dryspots - not visible in histological images, identified by low library sizes and no difference in mitochondrial ratio or 2) hangnail artefacts - library sizes and no difference in mitochondrial ratio.

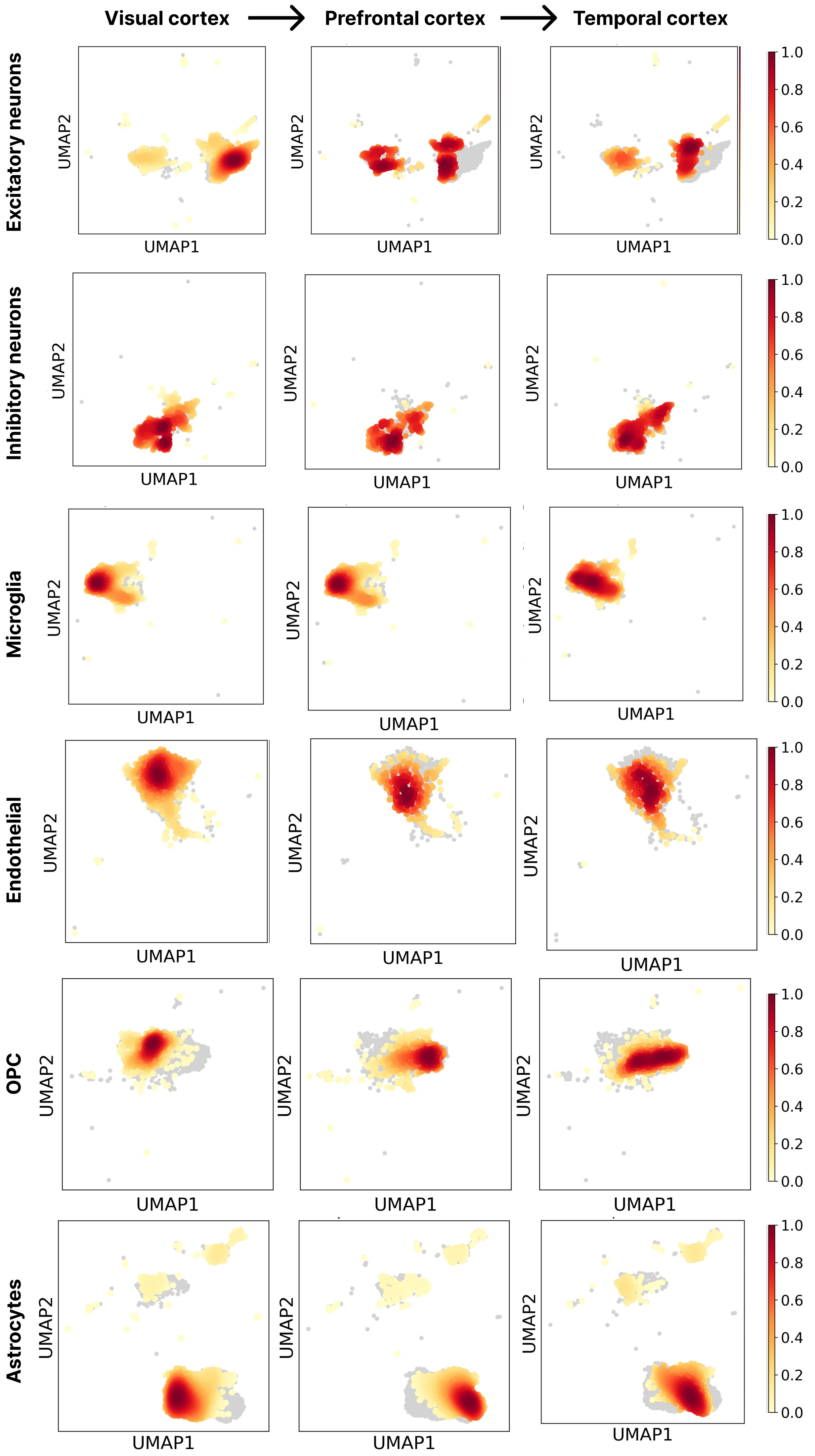

Visual cortex
Prefrontal cortex
Temporal cortex
Excitatory neurons
Inhibitory neurons
Microglia
Endothelial
OPC
Astrocytes
UMAP1
UMAP2
1.0
0.8
0.6
0.4
0.2
0.0

**Extended Data Figure 6: Regional density distributions per cell type mirror Alzheimer's disease progression gradient.**

UMAPs showing the kernel dentistry estimation (scanpy.tl.embedding_density) in all broad cell types (besides OL) across the temporal, prefrontal, and visual cortex. Regional gradients reflect decreasing AD pathology from lowest (visual cortex) to highest pathology (temporal cortex), based on pathological assumptions thereby providing a spatial snapshot of cellular changes along disease progression.

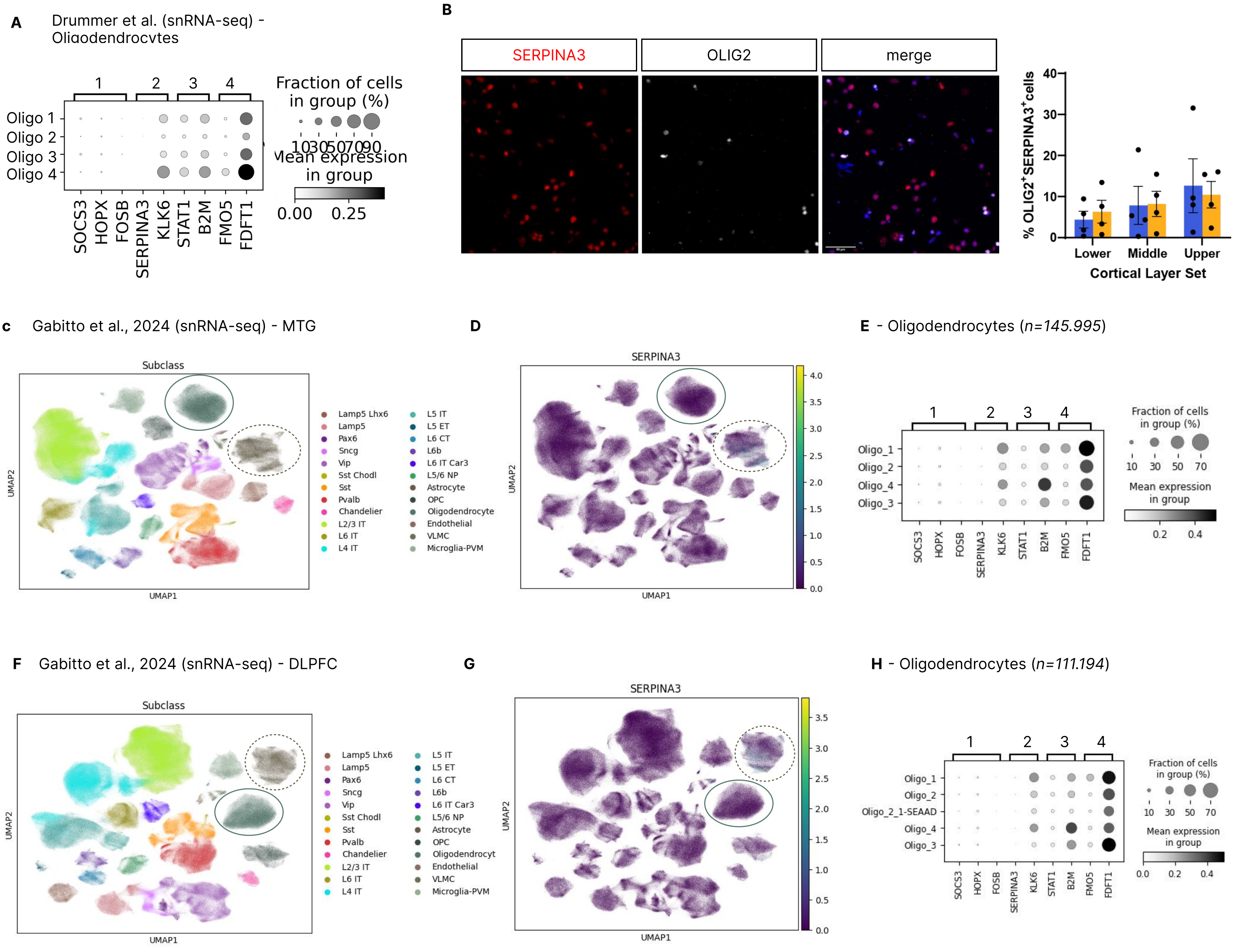


**Extended Data Figure 7: Common disease-associated oligodendrocyte markers are not expressed.**

**A** Dotplot showing the log-scales mean expression of common disease-associated OL (DAO) genes taken from the literature and divided into four groups: 1) transcription factors (TFs), 2) reactive markers, 3) immune-response genes and 4) genes associated with cholesterol metabolism. While TF expression might be lower in snRNA-seq vs scRNA-seq, other frequently cited DAO genes, such as SERPINA3, also do not show high expression. **B** Representative images and quantifications of immunohistochemical stainings of SERPINA3+OLIG2+ OL in human brains across cortical layers in the temporal cortex demonstrating a low abundance of SERPINA3+ OL and no specific enrichment in AD. Scale bars: 50 µm. **C-E** Analysis of SERPINA3 expression in the temporal cortex; snRNA-seq data from Gabitto et al., 2025. UMAP of all cell types (OL solid circle, astrocytes dashed circle) (**C**). UMAP overlay of SERPINA3 expression across celltypes, with the highest enrichment observed in astrocytes rather than oligodendrocytes (**D**). Dotplot showing commonly reported DAO genes in OL, supporting the observation that SERPINA3 is not highly expressed in disease OL (**E**). **F-H** Parallel analysis of SERPINA3 expression in dorsolateral prefrontal cortex (DLPFC); snRNA-seq data from Gabitto et al., 2025. UMAP of all cell types (OL solid circle, astrocytes dashed circle) (**F**). UMAP overlay of SERPINA3 expression across cell types, with the highest enrichment observed in astrocytes rather than oligodendrocytes, consistent with temporal cortex findings (**G**). Dotplot showing oligodendrocyte-specific marker gene expression in DLPFC, confirming the absence of SERPINA3 expression in oligodendrocytes across different cortical regions (**H**).

**Extended Table 1**

| Running numbe | Case ID | Brain area | Age | Sex | Disease Conditi | Braak Stage | APOE Genotype | Brainbank sourc | Experimental N | Sequencing batch |
|---|---|---|---|---|---|---|---|---|---|---|
| 1 | AD3 | Area Striata | 70 | male | AD | 6 | APOE 3/3 | NBB Munich | SJ_18 | 2 |
| 2 | AD3 | Temporal Gyrus | 70 | male | AD | 6 | APOE 3/3 | NBB Munich | SJ_23 | 2 |
| 3 | AD3 | Gyrus Rectus | 70 | male | AD | 6 | APOE 3/3 | NBB Munich | SJ_1 | 1 |
| 4 | AD5 | Area Striata | 74 | male | AD | 6 | APOE 3/3 | NBB Munich | SJ_25 | 2 |
| 5 | AD5 | Temporal Gyrus | 74 | male | AD | 6 | APOE 3/3 | NBB Munich | SJ_9 | 4 |
| 6 | AD5 | Gyrus Rectus | 74 | male | AD | 6 | APOE 3/3 | NBB Munich | SJ_19 | 4 |
| 7 | AD6 | Area Striata | 77 | male | AD | 5 | APOE 3/3 | NBB Munich | SJ_20 | 2 |
| 8 | AD6 | Temporal Gyrus | 77 | male | AD | 5 | APOE 3/3 | NBB Munich | SJ_8 | 1 |
| 9 | AD6 | Gyrus Rectus | 77 | male | AD | 5 | APOE 3/3 | NBB Munich | SJ_6 | 3 |
| 10 | AD7 | Area Striata | 75 | male | AD | 5 | APOE 3/3 | NBB Munich | SJ_24 | 3 |
| 11 | AD7 | Temporal Gyrus | 75 | male | AD | 5 | APOE 3/3 | NBB Munich | SJ_15 | 1 |
| 12 | AD7 | Gyrus Rectus | 75 | male | AD | 5 | APOE 3/3 | NBB Munich | SJ_34 | 3 |
| 13 | Ctrl3 | Area Striata | 70 | male | Ctrl | 1 | APOE 3/3 | NBB Munich | SJ_11 | 4 |
| 14 | Ctrl3 | Temporal Gyrus | 70 | male | Ctrl | 1 | APOE 3/3 | NBB Munich | SJ_17 | 1 |
| 15 | Ctrl3 | Gyrus Rectus | 70 | male | Ctrl | 1 | APOE 3/3 | NBB Munich | SJ_14 | 1 |
| 16 | Ctrl4 | Area Striata | 76 | male | Ctrl | 1 | APOE 3/3 | NBB Munich | SJ_33 | 2 |
| 17 | Ctrl4 | Temporal Gyrus | 76 | male | Ctrl | 1 | APOE 3/3 | NBB Munich | SJ_22 | 3 |
| 18 | Ctrl4 | Gyrus Rectus | 76 | male | Ctrl | 1 | APOE 3/3 | NBB Munich | SJ_5 | 4 |
| 19 | 20130894 | BA17 | 80 | male | Ctrl | 2 | APOE 3/3 | Newcastle Brain | SJ_16 | 2 |
| 20 | 20130894 | BA20/21 | 80 | male | Ctrl | 2 | APOE 3/3 | Newcastle Brain | SJ_37 | 1 |
| 21 | 20130894 | BA11 | 80 | male | Ctrl | 2 | APOE 3/3 | Newcastle Brain | SJ_26 | 3 |
| 22 | 20110891 | BA17 | 73 | male | Ctrl | 0 | APOE 3/3 | Newcastle Brain | SJ_32 | 1 |
| 23 | 20110891 | BA20/21 | 73 | male | Ctrl | 0 | APOE 3/3 | Newcastle Brain | SJ_30 | 2 |
| 24 | 20110891 | BA11 | 73 | male | Ctrl | 0 | APOE 3/3 | Newcastle Brain | SJ_10 | 3 |
| 25 | 20100742 | BA17 | 67 | male | Ctrl | 0 | APOE 3/3 | Newcastle Brain | SJ_27 | 2 |
| 26 | 20100742 | BA20/21 | 67 | male | Ctrl | 0 | APOE 3/3 | Newcastle Brain | SJ_29 | 4 |
| 27 | 20100742 | BA11 | 67 | male | Ctrl | 0 | APOE 3/3 | Newcastle Brain | SJ_12 | 3 |
| 28 | 20185928 | BA17 | 69 | male | AD | 6 | APOE 3/3 | Newcastle Brain | SJ_7 | 4 |
| 29 | 20185928 | BA20/21 | 69 | male | AD | 6 | APOE 3/3 | Newcastle Brain | SJ_4 | 2 |
| 30 | 20185928 | BA11 | 69 | male | AD | 6 | APOE 3/3 | Newcastle Brain | SJ_38 | 3 |
| 31 | 20174945 | BA17 | 69 | male | AD | 6 | APOE 3/3 | Newcastle Brain | SJ_21 | 4 |
| 32 | 20174945 | BA20/21 | 69 | male | AD | 6 | APOE 3/3 | Newcastle Brain | SJ_36 | 1 |
| 33 | 20174945 | BA11 | 69 | male | AD | 6 | APOE 3/3 | Newcastle Brain | SJ_13 | 4 |
| 37 | SD030/18 | BA17 | 63 | male | Ctrl | 0 | APOE 3/3 | Edinburgh Brain | SJ_2 and SJ_35 | 1 |
| 38 | SD030/18 | BA20/21 | 63 | male | Ctrl | 0 | APOE 3/3 | Edinburgh Brain | SJ_31 | 4 |
| 39 | SD030/18 | BA11/12 | 63 | male | Ctrl | 0 | APOE 3/3 | Edinburgh Brain | SJ_28 | 1 |
| 42 | SD006/14 | BA11/12 | 60 | male | Ctrl | 0 | APOE 3/3 | Edinburgh Brain | SJ_3 | 3 |

**Extended Table 2**

| Patient/Case ID | Cortical Area | Sample ID | Nr. cells | Mean reads per cell | Median genes per cell |
|---|---|---|---|---|---|
| | | | Cellranger output | | |
| | AS / VC | 22Jan4-DL008 | 1591 | 170.194 | 3.653 |
| Ctrl 1 (20100742) | GR / PFC | 22Jan4-DL004 | 583 | 836.419 | 4.339 |
| | TG / TC | 22Jan6-DL005 | 588 | 1.106.533 | 3.448 |
| | AS / VC | 22Jan3-DL006 | 1548 | 658.341 | 3.615 |
| Ctrl 2 (20110891) | GR / PFC | 22Jan4-DL003 | 1005 | 939.004 | 3.223 |
| | TG / TC | 22Jan4-DL007 | 1207 | 1.032.852 | 2.823 |
| | AS / VC | 22Jan6-DL003 | 1391 | 462.455 | 1.583 |
| Ctrl3 | GR / PFC | 22Jan3-DL004 | 428 | 1.171.714 | 3.786 |
| | TG / TC | 22Jan3-DL003 | 541 | 507.150 | 3.501 |
| | AS / VC | 22Jan4-DL005 | 6402 | 229.654 | 561 |
| Ctrl4 | GR / PFC | 22Jan6-DL004 | 1099 | 200.416 | 2.481 |
| | TG / TC | 22Jan5-DL005 | 1490 | 869.753 | 2.629 |
| | AS / VC | 22Jan4-DL006 | 434 | 549.543 | 1.460 |
| Ctrl 5 (20130894) | GR / PFC | 22Jan5-DL006 | 1055 | 49.015 | 3.626 |
| | TG / TC | 22Jan3-DL005 | 767 | 1.055.981 | 3.493 |
| | AS / VC | | | | |
| Ctrl 6 (SD00614) | GR / PFC | 22Jan5-DL010 | 510 | 257.023 | 238 |
| | TG / TC | | | | |
| | AS / VC | 22Jan3-DL008 | 1023 | 702.608 | 3.186 |
| Ctrl 7 (SD03018) | GR / PFC | 22Jan3-DL009 | 851 | 571.050 | 2.558 |
| | TG / TC | 22Jan6-DL009 | 2004 | 594.038 | 616 |
| | AS / VC | 22Jan6-DL007 | 1569 | 561.319 | 1.457 |
| AD1 (20174945) | GR / PFC | 22Jan6-DL008 | 774 | 313.528 | 2.861 |
| | TG / TC | 22Jan3-DL007 | 1210 | 960.979 | 1.789 |
| | AS / VC | 22Jan6-DL006 | 930 | 372.966 | 916 |
| AD 2 (20185928) | GR / PFC | 22Jan5-DL009 | 1185 | 432.585 | 3.493 |
| | TG / TC | 22Jan4-DL009 | 1215 | 218.908 | 827 |
| | AS / VC | 22Jan4-DL001 | 444 | 622.869 | 856 |
| AD3 | GR / PFC | 22Jan5-DL001 | 1433 | 136.383 | 262 |
| | TG / TC | 22Jan4-DL002 | 1079 | 236.137 | 1.145 |
| | AS / VC | 22Jan5-DL007 | 1163 | 129.906 | 391 |
| AD5 | GR / PFC | 22Jan6-DL002 | 186 | 358.891 | 1.040 |
| | TG / TC | 22Jan6-DL001 | 709 | 173.991 | 819 |
| | AS / VC | 22Jan5-DL008 | 1772 | 411.751 | 1.328 |
| AD6 | GR / PFC | 22Jan5-DL002 | 1124 | 472.439 | 2.601 |
| | TG / TC | 22Jan3-DL001 | 753 | 502.299 | 1.018 |
| | AS / VC | 22Jan5-DL003 | 914 | 171.668 | 2.415 |
| AD7 | GR / PFC | 22Jan5-DL004 | 994 | 332.127 | 3.088 |
| | TG / TC | 22Jan3-DL002 | 569 | 495.852 | 2.904 |

**Extended Table 3**

| | Nr. cells | Endothelial | Microglia-PVM | Neuronal: Gluta | Neuronal: GABA | Oligodendrocyte | OPC | Astrocyte | VLMC | Thresholding |
|---|---|---|---|---|---|---|---|---|---|---|
| 0 | 3492 | 0,000562 | 0,035383 | 0,76355 | 0,021904 | 0,15754 | 0,010952 | 0,004212 | 0,005897 | >0.75 |
| 1 | 2516 | 0,000393 | 0 | 0 | 0 | 0,999607 | 0 | 0 | 0 | >0.4 |
| 10 | 525 | 0 | 0,003025 | 0,000605 | 0,001815 | 0 | 0,991531 | 0,000605 | 0,00242 | >0.1 |
| 11 | 1176 | 0,956487 | 0,007911 | 0,002373 | 0,001582 | 0,000791 | 0 | 0 | 0,030854 | |
| 12 | 979 | 0 | 0,005547 | 0,027734 | 0,961173 | 0,002377 | 0 | 0,002377 | 0,000792 | |
| 13 | 940 | 0,000831 | 0 | 0,027431 | 0 | 0,004988 | 0 | 0,96675 | 0 | |
| 14 | 712 | 0,001043 | 0,008342 | 0,023983 | 0,965589 | 0 | 0,001043 | 0 | 0 | |
| 15 | 896 | 0 | 0,006494 | 0,002165 | 0,001082 | 0,001082 | 0 | 0,984848 | 0,004329 | |
| 16 | 696 | 0 | 0 | 0 | 0 | 0,998584 | 0 | 0 | 0,001416 | |
| 17 | 592 | 0,00146 | 0 | 0 | 0 | 0,994161 | 0,00146 | 0,00146 | 0,00146 | |
| 18 | 618 | 0,00319 | 0,001595 | 0 | 0 | 0,99362 | 0 | 0,001595 | 0 | |
| 19 | 550 | 0 | 0 | 0,006745 | 0 | 0 | 0 | 0,993255 | 0 | |
| 2 | 1929 | 0,001246 | 0,003739 | 0,989614 | 0,004155 | 0 | 0 | 0,000415 | 0,000831 | |
| 20 | 406 | 0,009141 | 0,117002 | 0,404022 | 0,027422 | 0,182815 | 0,04936 | 0,202925 | 0,007313 | |
| 21 | 296 | 0,001898 | 0,106262 | 0,189753 | 0,02277 | 0,056926 | 0,616698 | 0,003795 | 0,001898 | |
| 22 | 481 | 0,007722 | 0,009653 | 0,009653 | 0 | 0 | 0 | 0 | 0,972973 | |
| 23 | 146 | 0 | 0 | 0,00237 | 0 | 0 | 0,99763 | 0 | 0 | |
| 24 | 269 | 0 | 0 | 0 | 0,003425 | 0 | 0 | 0,989726 | 0,006849 | |
| 25 | 153 | 0,005405 | 0,2 | 0,275676 | 0,037838 | 0,12973 | 0,075676 | 0,2 | 0,075676 | |
| 26 | 39 | 0 | 0,981132 | 0 | 0 | 0,018868 | 0 | 0 | 0 | |
| 27 | 8 | 0 | 0,435897 | 0,179487 | 0,025641 | 0,102564 | 0 | 0,25641 | 0 | |
| 3 | 1754 | 0,000426 | 0,008514 | 0,007237 | 0,000851 | 0,982546 | 0,000426 | 0 | 0 | |
| 4 | 2065 | 0,001358 | 0,99321 | 0,002263 | 0 | 0,000905 | 0 | 0,000905 | 0,001358 | |
| 5 | 2078 | 0,000465 | 0 | 0,00093 | 0 | 0,998605 | 0 | 0 | 0 | |
| 6 | 2041 | 0,002395 | 0,194444 | 0,547414 | 0,037356 | 0,153736 | 0,02251 | 0,03592 | 0,006226 | |
| 7 | 1966 | 0 | 0,001514 | 0 | 0 | 0 | 0 | 0,998486 | 0 | |
| 8 | 1895 | 0,004644 | 0,459752 | 0,241486 | 0,089267 | 0,190402 | 0,001548 | 0,009804 | 0,003096 | |
| 9 | 1591 | 0,000604 | 0,006643 | 0,012681 | 0,000604 | 0,007246 | 0 | 0,969807 | 0,002415 | |

mixed-lineage cluster

low cell count clusters

**Extended Table 4**

| | Cell_Type | Marker_Genes | N_Genes | N_Samples | Pearson_r | Pearson_p | Spearman_r | Spearman_p |
|---|---|---|---|---|---|---|---|---|
| 0 | Astrocytes | SLC1A2, GFAP, AQ | 3 | 35 | -0,559913427 | 0,000469688 | -0,509426708 | 0,001774252 |
| 1 | Endo | BSG, CLDN5, FLT1 | 3 | 35 | -0,35764246 | 0,034916122 | -0,390560476 | 0,020359442 |
| 2 | Excitatory neurons | GRIN2A, CUX2, SLC | 4 | 35 | -0,689056766 | 4,72E-06 | -0,707537095 | 2,01E-06 |
| 3 | Inhibitory neurons | GAD2, GAD1, NXPH | 3 | 35 | -0,671220199 | 1,02E-05 | -0,713197391 | 1,52E-06 |
| 4 | Microglia | APBB1IP, C3, P2RY | 3 | 34 | -0,527473038 | 0,001347606 | -0,48650623 | 0,003528307 |
| 5 | OPC | OLIG1, PDGFRA, V | 4 | 35 | -0,424912729 | 0,010946942 | -0,469804631 | 0,004406051 |
| 6 | Oligodendrocytes | MOBP, MBP, PLP1 | 3 | 35 | -0,403558181 | 0,016216031 | -0,384900179 | 0,022420904 |
| 7 | VLMC | COL1A1, COL1A2 | 2 | 35 | -0,06808003 | 0,697578822 | -0,266089855 | 0,122335956 |

**Extended Table 5**

| Condition | Area | Threshold | | | | | |
|---|---|---|---|---|---|---|---|
| | | Nr. counts | | Gene counts | | MT | log10 MALAT1 |
| | | Min | Max | Min | Max | | |
| Control | PFC | 500 | 60,000 | 410 | 10,000 | 0,1 | 1 |
| | TC | 500 | 60,000 | 410 | 9,500 | 0,1 | 0,8 |
| | VC | 500 | 60,000 | 400 | 10,000 | 0,17 | 0,8 |
| AD | PFC | 500 | 50,000 | 400 | 9,000 | 0,1 | 1 |
| | TC | 500 | 30,000 | 350 | 7,000 | 0,2 | 0,8 |
| | VC | 500 | 40,000 | 350 | 8,000 | 0,25 | 0,8 |
